# A virus that has gone viral: Amino acid mutation in S protein of Indian isolate of Coronavirus COVID-19 might impact receptor binding and thus infectivity

**DOI:** 10.1101/2020.04.07.029132

**Authors:** Priyanka Saha, Arup Kumar Banerjee, Prem Prakash Tripathi, Amit Kumar Srivastava, Upasana Ray

## Abstract

Since 2002, beta coronaviruses (CoV) have caused three zoonotic outbreaks, SARS-CoV in 2002, MERS-CoV in 2012, and the recent outbreak of SARS-CoV-2 late in 2019 (also named as COVID-19 or novel coronavirus 2019 or nCoV2019. Spike(S) protein, one of the structural proteins of this virus plays key role in receptor (ACE2) binding and thus virus entry. Thus, this protein has attracted scientists for detailed study and therapeutic targeting. As the 2019 novel coronavirus takes its course throughout the world, more and more sequence analyses are been done and genome sequences getting deposited in various databases. From India two clinical isolates have been sequenced and the full genome deposited in GenBank. We have performed sequence analyses of the spike protein of the Indian isolates and compared with that of the Wuhan, China (where the outbreak was first reported). While all the sequences of Wuhan isolates are identical, we found point mutations in the Indian isolates. Out of the two isolates one was found to harbour a mutation in its Receptor binding domain (RBD) at position 407. At this site arginine (a positively charged amino acid) was replaced by isoleucine (a hydrophobic amino acid that is also a C-beta branched amino acid). This mutation has been seen to change the secondary structure of the protein at that region and this can potentially alter receptor ding of the virus. Although this finding needs further validation and more sequencing, the information might be useful in rational drug designing and vaccine engineering.

## Introduction

A virus gone viral. First case of COVID-19 was reported in December 2019 at Wuhan (China) and then it has spread worldwide becoming a pandemic, with maximum death cases in Italy, although initially, the maximum mortality was reported from China[1]. According to a WHO report, as of 02.04.2020 there were confirmed 8, 23,626 COVID-19 cases and 40598 deaths, that includes cases which were both locally transmitted or imported[2]. There are published reports which suggest that SARS-CoV2 shares highest similarity with bat SARS-CoV[3]. Scientists across the globe are trying to elucidate the genome characteristics using phylogenetic, structural and mutational studies[4]. Spike protein, one of the key proteins of SARS-CoV2 is involved directly with virus infection as it is involved in receptor recognition, attachment, binding and entry [5–7]. Sequence analyses of the spike protein can give us a plethora of information which can be instrumental in drug and vaccine development. In the present piece of work, we retrieved S protein sequences of the SARS-CoV2 from different geographical locations to identify notable features of S protein especially in Indian isolates. These analyses include identification of mutational signatures and their correlation with virus infection. Our analyses show unique point mutations in the spike protein of the Indian subtypes.

## Methods

Since COVID 19 or SARS-CoV-2 started from Wuhan, China, we started our analyses with Spike protein sequences from Wuhan. For our study we have considered all the full-length sequences that were available in GenBank. We first compared 17 available S protein sequences from Wuhan. Since they showed 100% sequence similarities, we considered one of these for our further analyses. Since Italy has also been affected aggressively by COVID-19, we included the sequence in our study. In this paper we have focussed on COVID-19 isolates from India. Till date only two complete genomes of Indian COVID-19 have been submitted in the database. For our sequence alignments we have used NCBI BLAST, CLUSTAL W and CLUSTAL OMEGA. To predict secondary structure, we have used CFSSP (Chou and Fasman secondary structure prediction) server.

Mutprep server was used to analyse the mutation. JMol and ConSurf tools were used to predict the structure of the proteins. PyMoL standalone software was used to visualize the structure and understand the pattern of bonding. Further kinetics and structure analyses were performed by the Dynemut Server and Chimera version 11.

## Results and Discussion

SARS-CoV-2 sequence data is expanding rapidly in the databases as the virus spreads worldwide. Although many sequences from various countries have been deposited, limited full genome sequences are available from most of the countries. This virus has infected people in various countries like China, Italy, Spain, USA, Germany, France, United Kingdom, India and many more and the data gets updated almost regularly by the World Health Organization (WHO). As of now, compared to many countries, the rate of transmission is comparatively controlled in India. Although this might be influenced by many factors like general immunity, point of entry of this virus in the country, measures taken to contain the spread, diagnosis, data management etc, we have used the available sequence data of Indian isolates to understand the biology of this virus.

From India, there are only two full genome sequences submitted from the state of Kerela (GenBank accession numbers MT012098 {(isolate SARS-CoV-2/human/IND/29/2020 or isolate 29) and MT050493 (isolate SARS-CoV-2/human/IND/166/2020 or isolate 166)}. We have compared the S protein sequence from these two isolates with that of Wuhan. All the 17 sequences from Wuhan that were first aligned to check sequence variability showed 100% sequence similarity (Figure 1).

**Figure 1:**
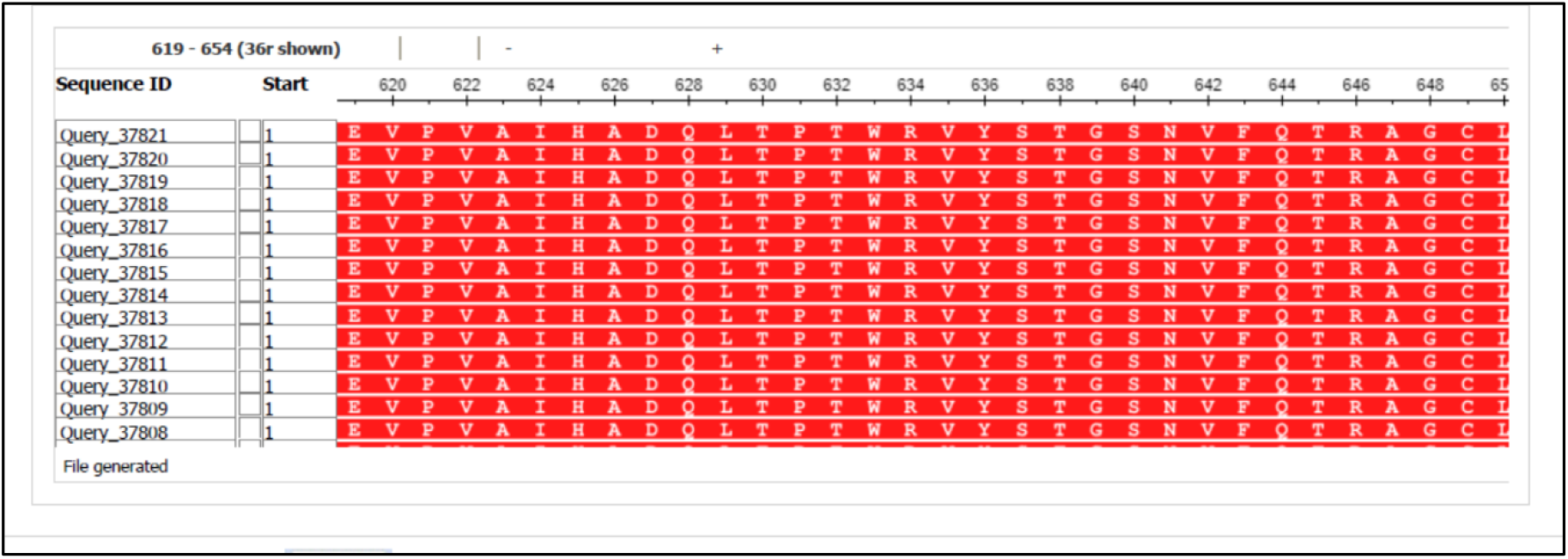
Multiple Sequence Alignment of Spike protein sequences of Wuhan isolates. 17 Spike protein sequences available in GenBank were aligned using NCBI BLASTp online tool and the multiple sequence alignment result has been shown.

To compare the Indian isolates, we aligned the S protein sequences of these isolates with Wuhan isolates and a sequence from Italy. While Wuhan and Italian isolates matched completely, we found few mutations in case of Indian isolates 29 and 166 as shown in Figure 2.

**Figure 2:**
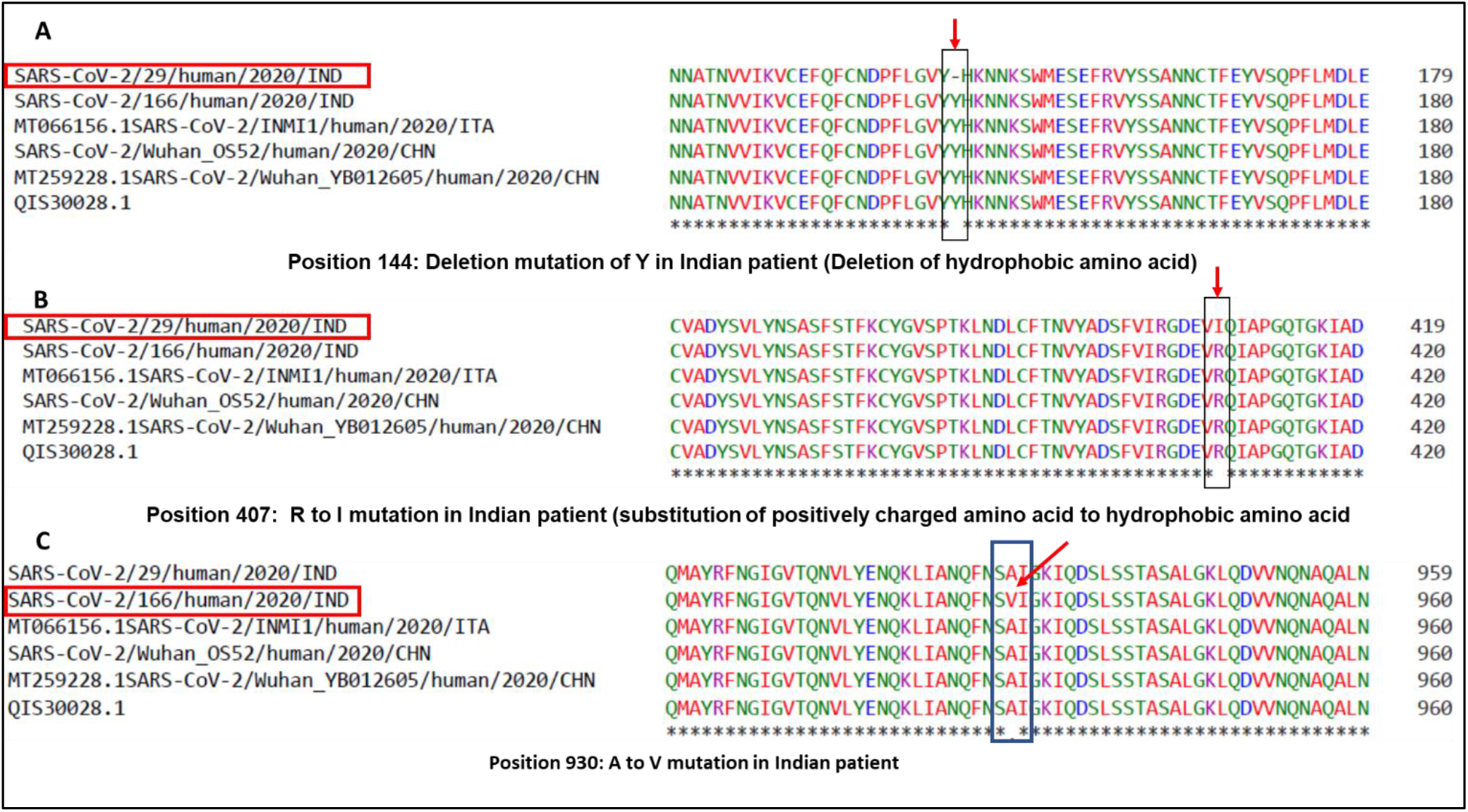
Multiple Sequence Alignment of Spike protein sequences of Indian, Wuhan and Italian isolates. Spike protein sequences of Indian Isolates available in GenBank were aligned using CLUSTAL Omega online tool with colour coding option and the multiple sequence alignment result has been shown above. The Indian isolates having mutations have been highlighted with red boxes. The mutations have been marked with red arrows. Panel A: Deletion mutation of Y in isolate 29. Panel B: Substitution mutation R->I at position 407 in isolate 29. Panel C: Substitution mutation of A->V at position 930 in isolate 166.

We observed that isolate 29 had two mutations, a deletion mutation where Y (tyrosine) at position 144 was absent as compared to Wuhan and Italian isolates. Due to deletion of the amino acid residues in position 144 in the protein structure there is a change in the beta sheets (Figure 3). This alteration may change the orientation of the molecule and also the stability of the protein itself. Ramachandran plot of this structure show slight shifts of angle in the β sheet configuration.

**Figure 3:**
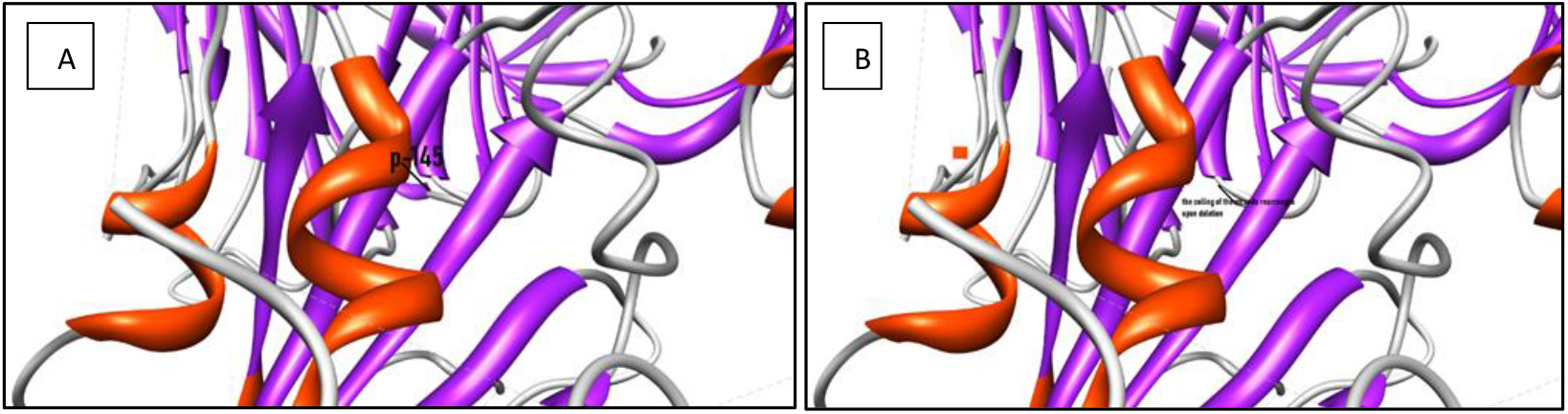
Deletion of the amino acid in 144 in Indian isolate 29. Panel A shows structure without deletion of amino acid tyrosine at position 144.Pnel B shows coiling of the strands rearranged after the deletion

On the other hand, at position 407, the same isolate had a substitution mutation of R (Arginine) to I (Isoleucine) (R407I). The receptor binding domain or RBD of the spike protein of SARS-CoV-2 lie between amino acids 331-524 [8]. Thus, the mutation R407I lie in the RBD which play key role in receptor binding. Arginine is a positively charged amino acid and Isoleucine is a hydrophobic amino acid with C-beta branch. While positively charged amino acid could be more exposed, hydrophobic amino acids secure themselves away from the outer aqueous environment. Since nature of these amino acids are so different, a substitution of this nature might change the conformation locally and can impose functional alterations i.e. with respect to receptor interaction. To confirm this theory further, we ran a secondary structure prediction using CFSSP server and found that while in case of Wuhan isolate, this region had helix only (H) (Figure 4 A), in case of R407I in Indian isolate there was introduction of sheets (E) (Figure 4B). This suggests that a change in secondary structure occurs in case of RBD of spike protein of isolate 29 of the 2019 novel Coronavirus of India. Tertiary structure analyses showed that there because of the mutation there is an introduction of additional oxygen molecule to the next residue. The protein stability score drops sharply (−4.08) and the thereby its electrostatic force. Such a condition makes the protein flexible and might affect interaction with the receptor. Alteration to the structure will cause shift in the hydrogen bonds and also the bond angle, two main pre-requisite for strong interaction with the receptor. The hydrogen potential tends to increase from 10 to 13.2 in case of mutation.

**Figure 4A:**
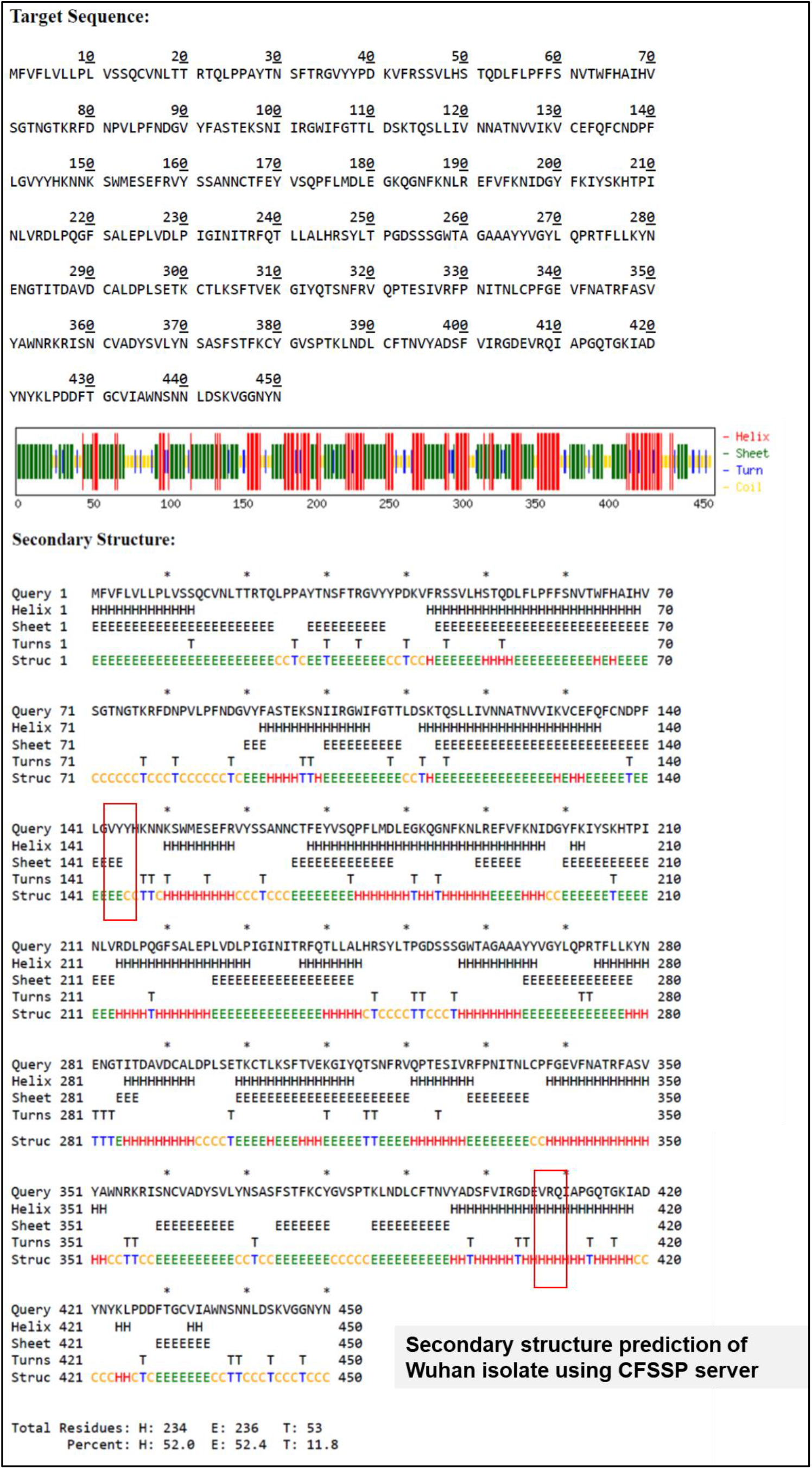
Secondary structure prediction of spike protein of Wuhan isolate. The area marked in red box has helix (H). This area has been seen to get mutated in Indian isolate

**Figure 4B:**
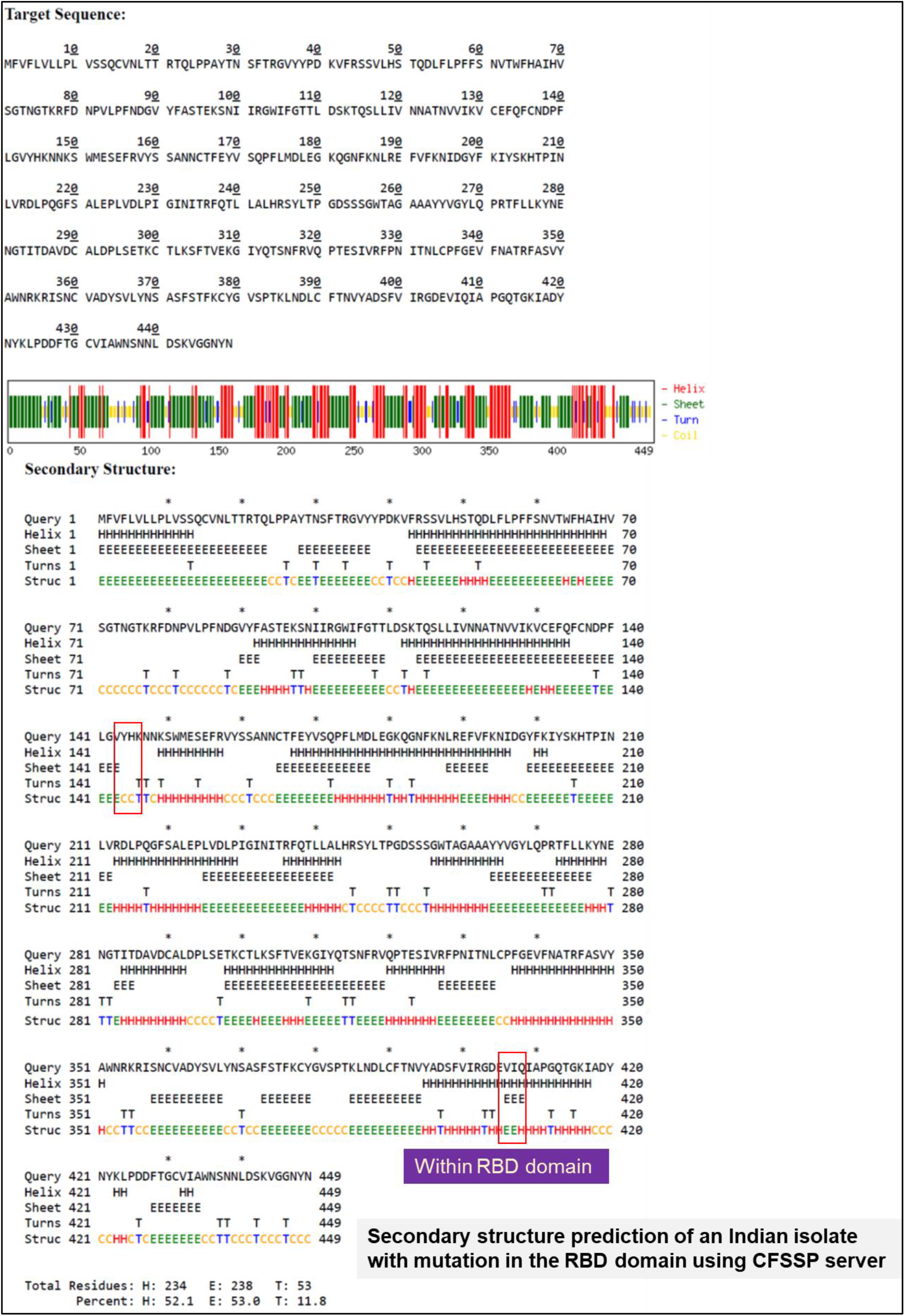
Secondary structure prediction of spike protein of Indian isolate 29. The area marked in red box that had helix (H) in Wuhan isolate shows introduction of sheets (E).

For the isolate 166 of India, we found a different mutation at position 930 of the spike protein (Figure 2C). Here there was a substitution of A (alanine) to V (valine) at position 930 (A930V). Since both the amino acids are hydrophobic in nature, any change that might occur due to this mutation might get masked upon tertiary structure formation and thus not imposing a functional change in the protein i.e. a conservative mutation. Despite this possibility, valine has some unique characteristics. Valine is one of the C-beta branched amino acids like threonine and isoleucine. C beta branched amino acids are bulkier towards the main chain and it is difficult for them to attain alpha helical conformations. Such amino acids have restricted conformations, are destabilizing in nature causing distortion in local helix backbone [9]. S protein if SARS-CoV2 has two domains: S1 and S2 [8]. While S1 has the RBD and in involved in receptor binding, S2 mediates fusion of viral and host cell membranes. Mutation A930V of Indian isolate 166 falls in the S2 subunit of S protein. Considering the nature of valine being destabilizing causing distortion, this mutation might have implications in viral membrane fusion subject to validation. Structure of the spike glycoprotein was retrieved from the protein data bank (PDB ID: 6VXX) (Figure 5). The residue alanine at position 930 though not associated with the active site of the molecule stabilizes majority of the chain A moiety in the protein due to its hydrophobic nature. On substitution with Valine in the same position, it can potentially change the affinity of the molecule towards its receptor.

**Figure 5:**
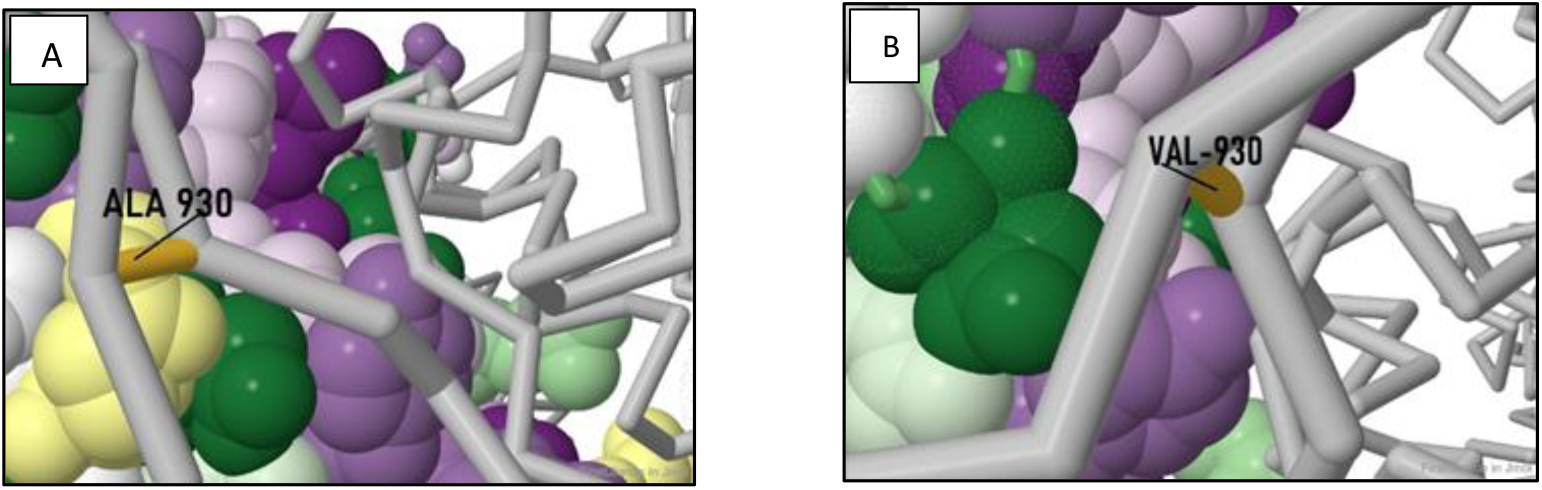
Alanine (Panel A) to Valine (Panel B) mutation in isolate 166.

Thus, taken together the mutations in S protein of Indian isolates can potentially alter virus entry and thus determine the infectivity of the virus. More sequence information along with mutational studies on receptor virus binding will further help strengthen this observation. The sites of mutation, geographic location, frequency of such mutations, knowledge about progression of infection and disease severities will help correlating the significance of these mutation with respect to virus evolution and virulence. Such information will further help in strategic designing of drug targets.

## Acknowledgement

We would like to thank Dr. Anupam Das Talukdar, Department of Life Sciences and Bioinformatics, Assam University for sharing server. We also thank Department of Biotechnology, DBT for funding to PS. CSIR is also acknowledged.

## References

[1] Guo Y.R., Cao Q.D., Hong Z.S., Tan Y.Y., Chen S.D., Jin H.J., Tan K.S., Wang D.Y., Yan Y., The origin, transmission and clinical therapies on coronavirus disease 2019 (COVID-19) outbreak–an update on the status, Military Medical Research, 7 (2020) 1–10.

[2] Ziff A.L., Ziff R.M., Fractal kinetics of Covid-19 pandemics (with update 3/1/20), MedRxiv preprint, (2020).

[3] Smith M., Smith J.C., Repurposing therapeutics for COVID-19: Supercomputer-based docking to the SARS-CoV-2 viral spike protein and viral spike protein-human ACE2 interface, (2020).

[4] Ahmed S.F., Quadeer A.A., McKay M.R., Preliminary identification of potential vaccine targets for the COVID-19 coronavirus (SARS-CoV-2) based on SARS-CoV immunological studies, Viruses, 12 (2020) 254.

[5] Wu Y., Strong evolutionary convergence of receptor-binding protein spike between COVID-19 and SARS-related coronaviruses, bioRxiv, (2020).

[6] Ortega J.T., Serrano M.L., Pujol F.H., Rangel H.R., Role of changes in SARS-CoV-2 spike protein in the interaction with the human ACE2 receptor: An in silico analysis, EXCLI journal, 19 (2020) 410.

[7] Z. Liu, X. Xiao, X. Wei, J. Li, J. Yang, H. Tan, J. Zhu, Q. Zhang, J. Wu, L. Liu, Composition and divergence of coronavirus spike proteins and host ACE2 receptors predict potential intermediate hosts of SARS‐CoV‐2, Journal of Medical Virology, (2020).

[8] Wanbo T., Lei H., Xiujuan Z., Jing P., Denis V., Shibo J., Characterization of the receptor-binding domain (RBD) of 2019 novel coronavirus: implication for development of RBD protein as a viral attachment inhibitor and vaccine, Cellular and Molecular Immunology, (2020).

[9] Cornish V.W., Kaplan M.I., Veenstra D.L., Kollman P.A., Schultz P.G. Stabilizing and Destabilizing Effects of Placing Beta-Branched Amino Acids in Protein Alpha-Helices

